# Gaming expertise induces meso-scale brain plasticity and efficiency mechanisms as revealed by whole-brain modeling

**DOI:** 10.1101/2023.08.21.554072

**Authors:** Carlos Coronel-Oliveros, Vicente Medel, Sebastián Orellana, Julio Rodiño, Fernando Lehue, Josephine Cruzat, Enzo Tagliazucchi, Aneta Brzezicka, Patricio Orio, Natalia Kowalczyk-Grębska, Agustín Ibáñez

## Abstract

Video games are a valuable tool for studying the effects of training and neural plasticity on the brain. However, the underlaying mechanisms related to plasticity-induced brain structural changes and their impact in brain dynamics are unknown. Here, we used a semi-empirical whole-brain model to study structural neural plasticity mechanisms linked to video game expertise. We hypothesized that video game expertise is associated with neural plasticity-mediated changes in structural connectivity that manifest at the meso-scale level, resulting in a more segregated functional network topology. To test this hypothesis, we combined structural connectivity data of StarCraft II video game players (VGPs, n = 31) and non-players (NVGPs, n = 31), with generic fMRI data from the Human Connectome Project and computational models, with the aim of generating simulated fMRI recordings. Graph theory analysis on simulated data was performed during both resting-state conditions and external stimulation. VGPs’ simulated functional connectivity was characterized by a meso-scale integration, with increased local connectivity in frontal, parietal and occipital brain regions. The same analyses at the level of structural connectivity showed no differences between VGPs and NVGPs. Regions that increased their connectivity strength in VGPs are known to be involved in cognitive processes crucial for task performance such as attention, reasoning, and inference. In-silico stimulation suggested that differences in FC between VGPs and NVGPs emerge in noisy contexts, specifically when the noisy level of stimulation is increased. This indicates that the connectomes of VGPs may facilitate the filtering of noise from stimuli. These structural alterations drive the meso-scale functional changes observed in individuals with gaming expertise. Overall, our work sheds light into the mechanisms underlying structural neural plasticity triggered by video game experiences.

## 1 Introduction

Video games have evolved from a mere form of entertainment to a prominent hallmark of modern culture. They have permeated several aspects of contemporary life, expanding beyond hobbies to include eSports, education, and medicine [1]. From a theoretical perspective, video games can be considered as enriched environments that provide engaging and ecological stimuli, motivating users to get involved in a wide range of activities. These environments can induce the coordination of different cognitive process such as attention, visuomotor skills, and reasoning [2]. The characteristics of an enriched environment, such as exploring innovative activities [3] and working under time pressure in a dynamic setting [4] are directly present in action video games [1,5,6] and support the notion that video games offer a naturalistic and valuable tool for studying how training and expertise induce structural and functional changes in the brain through neural plasticity [1,7].

Real-time strategy (RTS) video games have been shown to be a rich source of enjoyment, promoting learning and structural plasticity [5,8–11]. This genre encompasses a wide range of popular titles, with StarCraft II standing out as a successful game in both entertainment and professional eSports, offering over 200 million dollars prizes in world-class tournaments in 2022 (https://www.esportsearnings.com). The latter is a RTSs military science fiction video game developed by Blizzard Entertainment Inc. in 2010, requiring players to manage resources, build bases, create armies, and memorize keyboard shortcuts while engaging in combat with other players or computer artificial intelligent devices (**Fig 1**). The game demands significant cognitive efforts including reasoning, visual attentional skills, and inference abilities for making predictions. Previous studies have supported the use of RTS games for investigating neural plasticity mechanisms associated with expertise [8–13].

**Fig 1.**
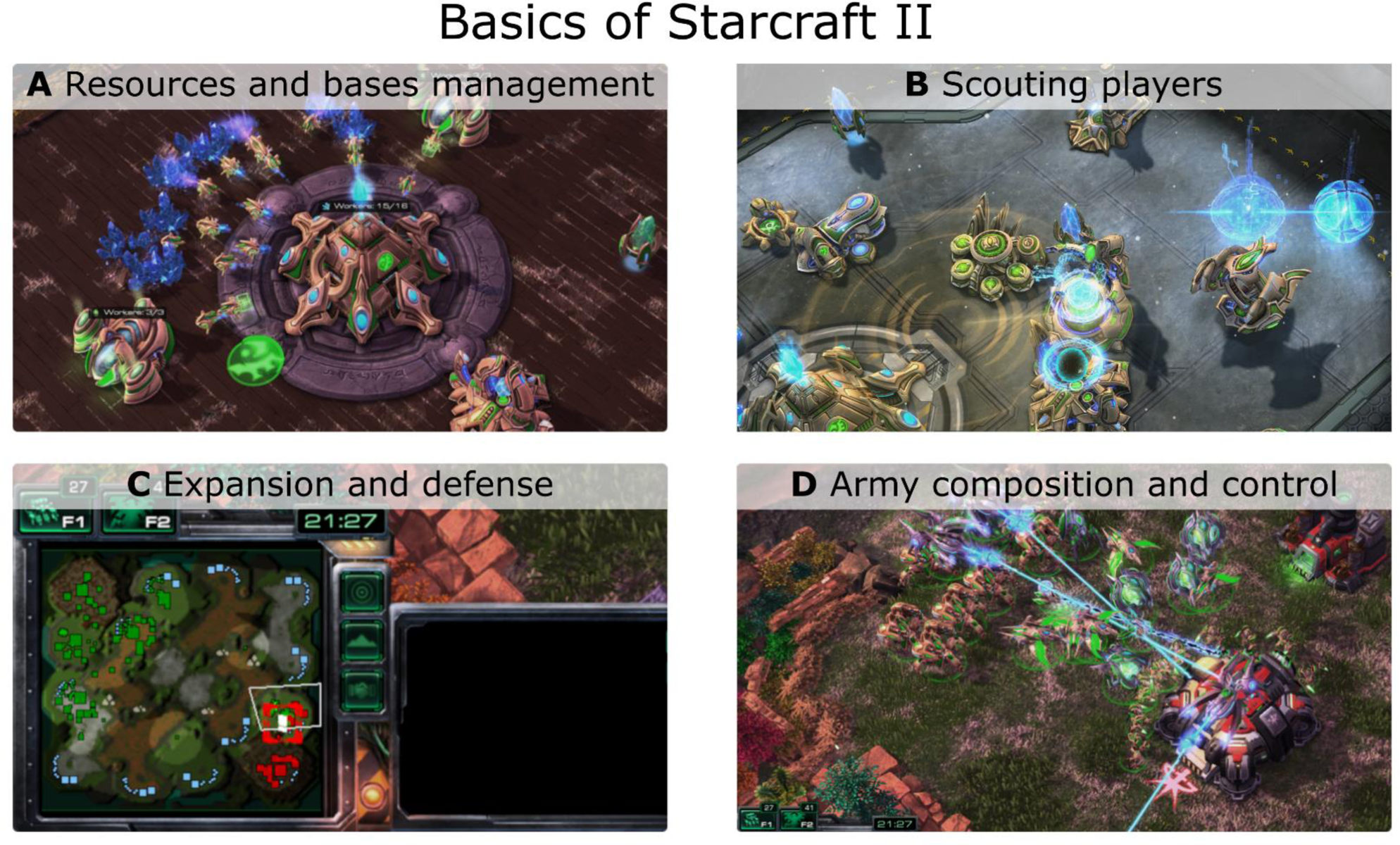
StarCraft II gameplay basics. The game requires that players multitask four principal aspects. A) Management of resources (minerals and “vespene gas”), create gatherers and builders, build new facilities and research technologies. B) Scout and spy other players, for example, using a “radar scan”, are essential actions to adapt own strategies and army composition in response to enemies’ actions. C) In the figure, the minimap shows the location of both players’ bases and units. Players should manage and defend multiple bases in the map. D) The game requires creating, managing, and controlling complex armies to fight with enemies and defend bases. The in-game screenshots of StarCraft II (©2023 Blizzard Entertainment, Inc.) were taken by the authors of this work.

In a previous work [9] we demonstrated that StarCraft II video game players (VGPs) exhibited increased total white matter connections between occipital and parietal areas and within occipital areas compared to non-video game players (NVGPs). Furthermore, VGPs showed altered structural local efficiency within the parieto-occipital subnetwork. The positive association between network metrics and game playing time suggests a close relationship between extensive, long-term RTS game play and neuroplastic changes [9]. Other studies have suggested that experiences with RTS games are associated with altered white matters networks connections in prefrontal, limbic, and sensorimotor networks [11], hippocampus and cingulum [10], and enhanced communication between dorsal attentional and salience networks [12]. Moreover, RTS and action video games have been linked to faster reaction times and improved visual attentional skills [1,5,6]. What is more, gray matter in brain regions such as the entorhinal cortex, hippocampus and occipital areas, which are mainly associated with memory [14] and visual attention [15], has been found to be related to the total amount of time spent playing video games [16]. These studies suggest that expertise and training in RTS games induce neural plasticity changes that may be associated with visual attentional performance. However, there is a gap in mechanistic explanations, particularly regarding how structural alterations driven by neural plasticity impact brain dynamics at the meso- and macro-scale levels. Similarly, how the specific brain dynamics associated with expertise can become more efficient [17–19], e. g., promoting a more expert early and late processing of visual information [20,21]. Whole-brain computational models provide a way to explore structure-function relationships [22,23] by generating brain activity based on anatomical connections and the dynamics of individual brain regions.

Whole-brain models unveils the biophysical principles underlying brain structure, function, and human cognition [22,23]. These computational models can be used to test mechanistic hypotheses and have been extensively used to characterize brain dynamics in health and disease [22–27], make predictions and recover latent information not directly available in real experimental setups [24], for data augmentation with the aim of training machine learning algorithms [25], and for data completion using computational simulations [26]. We used a semi-empirical whole-brain modeling approach to predict the potential functional connectivity (FC) differences between VGPs and NVGPs, guided by empirical structural connectivity (SC) data from VGPs and NVGPs [9]. The model, simulated fMRI BOLD signals and constructed FC matrices, and then we assessed the model fits by comparing simulated and empirical FCs. We exploited the capability of whole-brain modeling to generate missing data using data completion. By inferring the FC and responsivity to stimulation from players’ empirical anatomical connections, we were able to uncover latent information that was not available in the original experimental setup. This novel approach allowed us to provide a comprehensive characterization of the player’s brain dynamics, including both resting-state and in-silico stimulation conditions. Furthermore, our study employed a single-subject approach, surpassing generalizations beyond simple groups of participants, and integrated multimodal neuroimaging to gain a deeper understanding of the complex brain dynamics associated with expertise in video gaming. Notably, this is the first application of whole-brain modeling in the context of video game expertise.

Here, we analyzed brain FC in the context of video game expertise, by combining a whole-brain model, fMRI data from the Human Connectome Project (HCP) [27], and brain connectomes obtained from VGPs and NVGPs published in Kowalczyk-Grębska et al. [9]. We used a phenomenological model based on Stuart-Landau oscillators, able to reproduce whole-brain dynamics in diverse settings [22,28,29]. We simulated fMRI BOLD signals and constructed FC matrices and fitted the model to empirical data. Our first hypothesis posited that structural neural plasticity is the driving mechanism behind the meso-scale functional reorganization observed in the brains of VGPs [30], characterized by higher parieto-occipital and frontoparietal connectivity strength [9]. To test this hypothesis, we fitted the model to resting-state data and identified the key structural connections responsible for the alterations in functional connectivity among players. Our second hypothesis is that brain networks altered in VGPs are associated with specific cognitive skills linked to expertise in RTS games. We explored this hypothesis by analyzing association maps obtained using Neurosynth [31], allowing us to generalize our results from video games to other cognitive domains. Lastly, our final hypothesis proposed that VGPs’ connectomes exhibit a higher signal-to-noise ratio within the parieto-occipital loop [21,32,33], resulting in robustness to stimulation in noisy contexts because of videogame expertise [17]. We explored this hypothesis by applying in-silico external stimulation to homotopic pairs of occipital brain areas [34–36].

By simultaneously testing these hypotheses we showed that expertise-related changes in structural connectivity, induce a meso-scale functional re-organization of RTS players’ brain dynamics, changing the responsivity to external stimulation by tuning specific cognitive process.

## 2 Results

We built a whole-brain model (**Fig 2**) to predict video game players’ fMRI activity and FC from their brain anatomical connections, and to evaluate players responsivity to external stimulation. We informed the model with empirical priors (**Fig 2A**) which consisted of subject-specific empirical human structural connectivity matrices of VGPs and NVGPs. We used the connectomes reported in a structural network analysis of parieto-occipital connectivity in StarCraft II players [37] (VGPs = 31, NVGPs = 31 subjects). Our whole-brain model was used to infer FC of VGPs and NVGPs, not available in the original study [9], by data completion [38]. To ensure that the simulated FC patterns resembled actual empirical data, we informed the model with empirical fMRI BOLD FC data acquired from 100 subjects of the Human Connectome Project (HCP) Q3 release [39] (**Fig 2A**). We fitted each single-subject model to the grand-averaged FC matrix from HCP. Both the structural and functional data were parcellated using the automated anatomical labeling (AAL) brain parcellation [40] (**Fig 2B**, and regions listed in Table 1), keeping the 90 cortical and subcortical regions of interests (ROIs) (AAL90, **Table 1**). Each brain area in the model corresponds to one ROI of the AAL90 (circles on the brain of **Fig 2**). ROIs are connected using a structural connectivity matrix (SC), depicted as lines in the brain of **Fig 2B**. The local dynamics of each node were simulated using the Stuart-Landau oscillator [28,29] (**Fig 2B**) set close to the model bifurcation point, where noise-driven and sustained oscillations coexist in time [29,41]. Furthermore, the model contemplates noisy external stimulation –that can be turned “on” and “off”– applied to pairs of homotopic regions (the same ROI in both brain hemispheres) [28,42–44] (**Fig 2B**).

**Fig 2.**
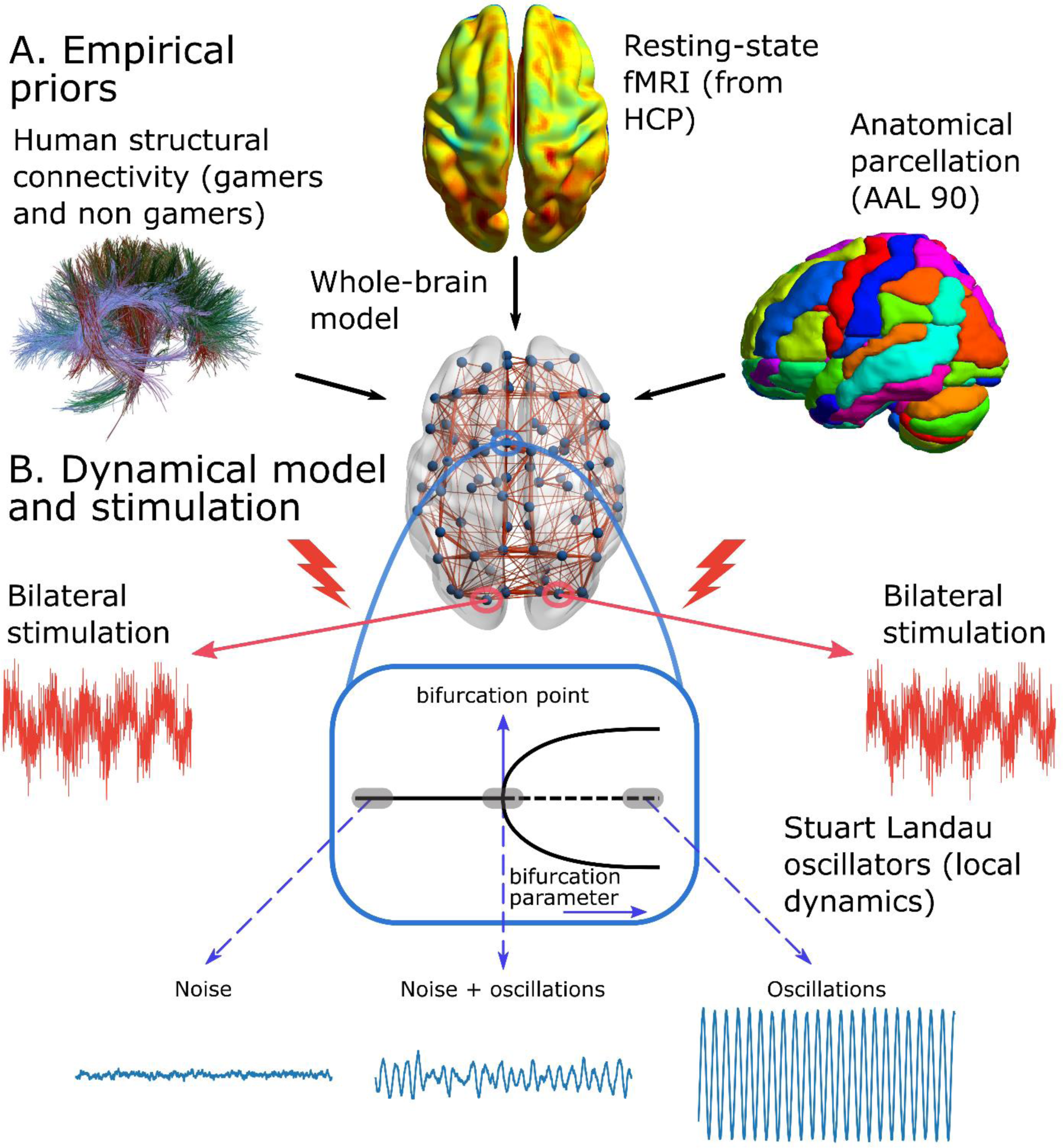
Whole-brain model. **A)** The computational model incorporates empirical priors using single-subject human connectomes obtained from players and non-players using DTI and functional connectivity data based on averaged fMRI BOLD signals from the Human Connectome Project (HCP). Both structural and functional connectivity data were parcellated using the automated anatomical labeling (AAL) atlas, which comprises 90 cortical and subcortical brain regions (AAL90). **B)** Brain regions (blue circles) are interconnected using the structural connectivity matrices and brain dynamics were modeled using the Stuart Landau oscillators. Within each area, a local bifurcation parameter controls the regional dynamics, which could exhibit noise-driven oscillations (at the left of the bifurcation point), sustained-oscillations (at the right) or a mixture of noise plus oscillations (close to the bifurcation point). By default, regional dynamics operates close to the Hopf bifurcation. Furthermore, the model contemplates homotopic stimulation, involving the application of periodic sinusoidal and noisy input to pair of homotopic brain areas (same region, on the left and right hemispheres).

**Table 1.**
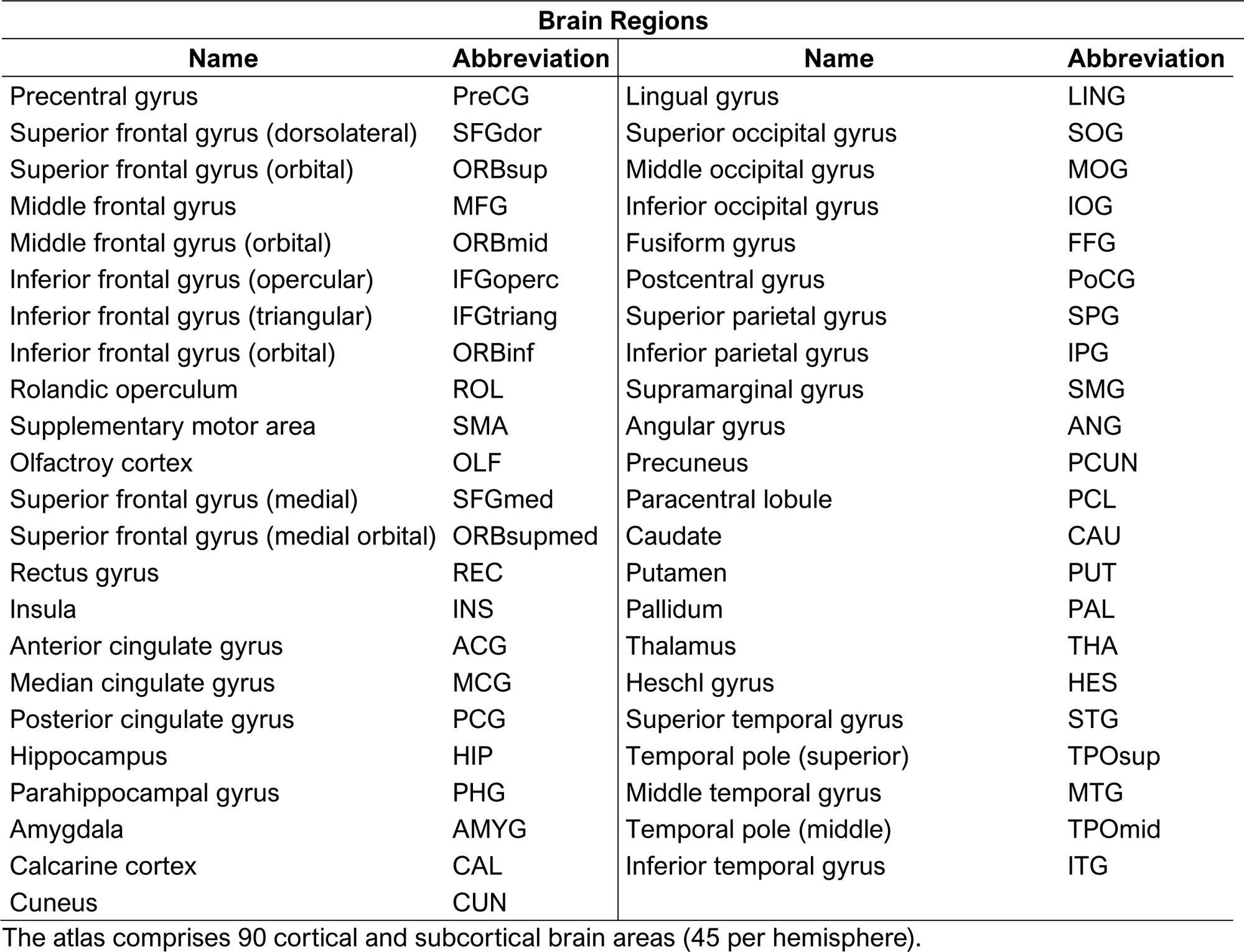
List of brain regions of AAL90 parcellation.

| Brain Regions |  |  |  |
| --- | --- | --- | --- |
| Name | Abbreviation | Name | Abbreviation |
| Precentral gyrus | PreCG | Lingual gyrus | LING |
| Superior frontal gyrus (dorsolateral) | SFGdor | Superior occipital gyrus | SOG |
| Superior frontal gyrus (orbital) | ORBsup | Middle occipital gyrus | MOG |
| Middle frontal gyrus | MFG | Inferior occipital gyrus | IOG |
| Middle frontal gyrus (orbital) | ORBmid | Fusiform gyrus | FFG |
| Inferior frontal gyrus (opercular) | IFGoperc | Postcentral gyrus | PoCG |
| Inferior frontal gyrus (triangular) | IFGtriang | Superior parietal gyrus | SPG |
| Inferior frontal gyrus (orbital) | ORBinf | Inferior parietal gyrus | IPG |
| Rolandic operculum | ROL | Supramarginal gyrus | SMG |
| Supplementary motor area | SMA | Angular gyrus | ANG |
| Olfactory cortex | OLF | Precuneus | PCUN |
| Superior frontal gyrus (medial) | SFGmed | Paracentral lobule | PCL |
| Superior frontal gyrus (medial orbital) | ORBsupmed | Caudate | CAU |
| Rectus gyrus | REC | Putamen | PUT |
| Insula | INS | Pallidum | PAL |
| Anterior cingulate gyrus | ACG | Thalamus | THA |
| Median cingulate gyrus | MCG | Heschl gyrus | HES |
| Posterior cingulate gyrus | PCG | Superior temporal gyrus | STG |
| Hippocampus | HIP | Temporal pole (superior) | TPOsup |
| Parahippocampal gyrus | PHG | Middle temporal gyrus | MTG |
| Amygdala | AMYG | Temporal pole (middle) | TPOmid |
| Calcarine cortex | CAL | Inferior temporal gyrus | ITG |
| Cuneus | CUN |  |  |

The atlas comprises 90 cortical and subcortical brain areas (45 per hemisphere).

The fitting procedure consisted in tuning the global coupling parameter, G, which scales the strength of the long-range connections, that is, the connections represented by the SC matrix (denoted by M in the model). For each subject within VGPs and NVGPs groups, we simulated BOLD-like signals (**Fig 3A**) for different values of G and model realizations (100 random seeds or replicates of the same simulations). Then, we computed the model’s FCs and compared it to the empirical FC matrices using the Structural Similarity Index (SSIM) [45] (**Fig 3A**). The SSIM constitutes the goodness-of-fit of the model fitting, with values closer to 1 indicating a good fit to empirical data. As an example, we show the fitting for one subject of VGPs and NVGPs in **Fig 3B**. The value of G that maximizes the SSIM was the best one for model fitting and was fixed across simulations. We found no differences, between VGPS and NVGPs, in G values (*t* = −1.858, *p* = 0.068) and SSIM (*t* = 0.464, *p* = 0.644) (**Fig 3B**). The mean SSIM values were 0.417 and 0.415, which are acceptable in the context of similar works in the field [44,46]. Thus, both VGPs and NVGPs models fitted equally good to the empirical HCP data. For visualization purposes, we show the empirical and simulated FC matrices in **Fig 3C**. Despite VGPs and NGVPs FCs looking similar, patterns of increased and decreased FC were observed when computing the difference matrix (**Fig 3C**).

**Fig 3.**
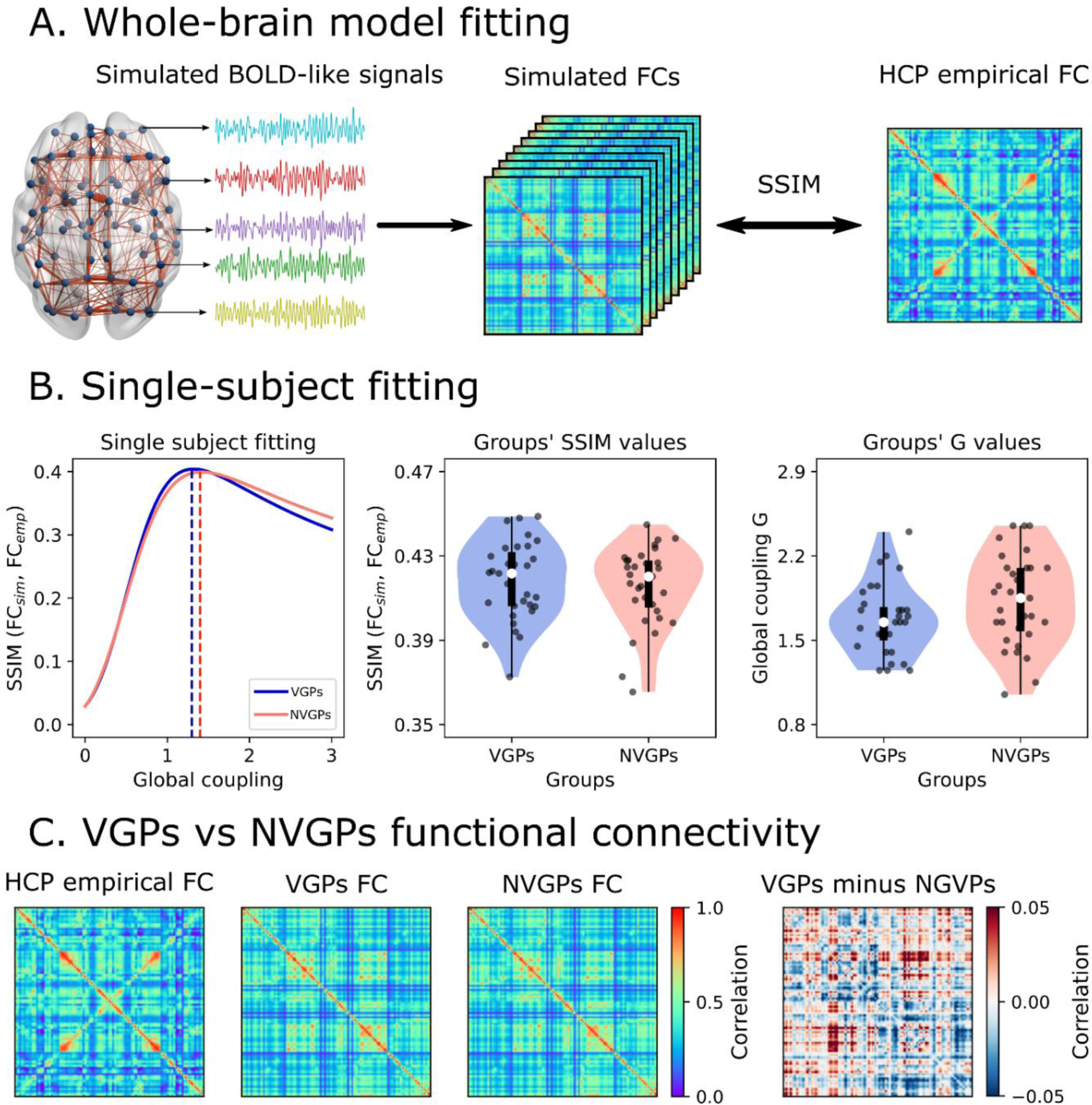
Single-subject fitting. **A)** The whole-brain model was used to simulate fMRI BOLD-like signals, which were then used to build functional connectivity (FC) matrices using pairwise Pearson’s correlation. We compared the simulated and empirical matrices using the SSIM, where values close to 1 indicate a good fit between the empirical (HCP) and modeled FC. **B)** We varied the global coupling parameter, *G*, and computed the structural similarity index (SSIM) across multiple model realizations using 100 random seeds) **T**he left panel shows an example for one subject each of video game players (VGPs) and non-video game players (NVGPs) conditions. The optimal *G* values were determined by maximizing the SSIM. The model was fitted to the data for both VGPs and NVGPs players across all subjects. Points in the violin plots represent individual subjects. **C)** Empirical (HCP) and simulated FC matrices (averaged across subjects). While VGPs and NVGPs matrices may appear similar, the difference matrix reveals distinctive patterns of increased and decreased functional connectivity in VGPs, as shown in the right panel. The data points in the violin plots correspond to subjects (VGPs = 31, NVGPs = 31). Box plots were built using the 1st and 3rd quartiles, the median as well as the maximum and minimum values of the distributions.

From the single-subject models, we continue to characterize the differences in FC between players and non-players in the next subsections.

### 2.1 Gaming expertise induces functional meso-scale integration

We first analyzed the differences between VGPs and NVGP in the FCs that arise from the different SCs using the fitted whole-brain model. FC connectivity strength was analyzed at the whole-brain level, and between parietal and occipital subnetworks. Further, graph theory tools were used to characterized topological properties of brain functional networks. Specifically, we used global efficiency as a measure of integration [47]. The metric is based on the concept of shortest paths; values of global efficiency close to 1 are a signature of a highly integrated brain network, where brain areas can easily reach each other towards multiple paths. On the other hand, transitivity was used to compute functional segregation [48]. It is a variant of the clustering coefficient and is based on the count of the triangular motifs of network. Values close to 1 indicate a more locally connected, that is, segregated network topology. We did not find any difference at the level of overall FC (global correlations) between VGPs and NVGPs (*t* = −1.471, *p* = 0.146, **Fig 4A**). Interestingly, parieto-occipital FC strength was higher in VGPs (*t* = 2.789, *p* = 0.007, **Fig 4A**), reflecting that those local differences are independent of the mean global correlations. Thus, these differences can be ascribed to the topological properties of players’ connectomes. When analyzing whole-brain functional network topology, we observed that VGPs’ FCs show a shift towards functional segregation involving higher transitivity (*t* = 2.705, *p* = 0.009) and lower global efficiency (*t* = −2.960, *p* = 0.004) in VGPs than NVGPs (**Fig 4A**). Correlations between the playing time (logarithm of the hours per week) and the FC metrics were null at overall FC (*r* = 0.204, *p* = 0.269), parieto-occipital FC (*r* = 0.196, *p* = 0.269) and integration (*r* = −0.285, *p* = 0.120) levels. A positive correlation with segregation (*r* = 0.356, *p* = 0.049) was observed. This suggests that a shift to more local-organized functional connectivity pattern is associated with VGPs, scaling with time spent playing video games, and suggesting a functional re-organization of brain networks as consequence of expertise.

**Fig 4.**
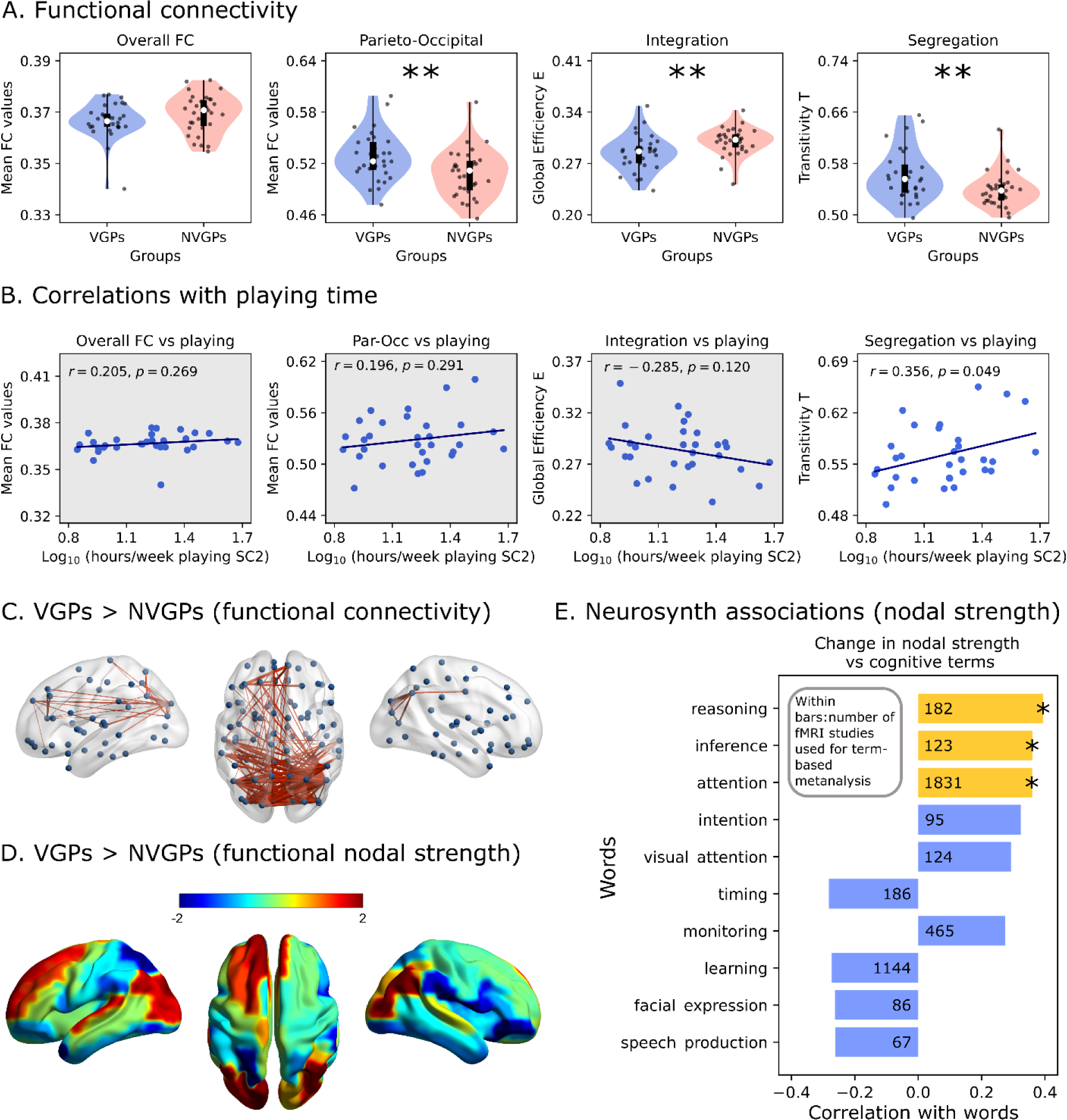
Resting-state functional connectivity of players and non-players. **A)** Global functional connectivity (FC), parieto-occipital FC, integration (global efficiency) and segregation (transitivity) of video game (VGPs) and non-video game (NVGPs) players during resting-state (without stimulation). **B)** Correlation between functional connectivity and playing time (logarithm of hours per week). **C)** Strengthened functional connections in VGPs compared to NVGPs, obtained using network-based statistics. **D)** Differences in regional functional nodal strength between VGPs and NVGPs. **E)** Correlations between the vector in D) and association maps, derived from Neurosynth [49,50], ascribed to 89 irreducible cognitive terms [51]. Correlations were FDR-corrected, and statistical significance is highlighted by yellow bars with stars. Data points represent individual subjects (VGPs = 31, NVGPs = 31). Box plots were built using the 1st and 3rd quartiles, the median, the maximum and minimum values of distributions. All correlations were computed using Pearson’s correlation coefficient. ***: *p* < 0.001, **: *p* < 0.01, *: *p* < 0.05. The number displayed within the bars corresponds to the number of fMRI studies used for the term-based meta-analysis. FDR corrected *p*-values were used.

Next, to gain insights about the specific FC patterns ascribed to VGPs, we used a Network-Based Statistics (NBS) analysis [52]. This method has demonstrated to have higher power, avoiding the multiple comparison problem [52]. We found a subnetwork that is strongly connected in VGPs than in NVGPs (*p*

< 0.001 using 10000 permutations, **Fig 4C**). Higher connectivity strength within the parieto-occipital loop is observed in VGPs. Some connections projecting from the posterior brain areas to the more frontal ones showed increased connectivity strength, suggesting that both parieto-occipital and frontoparietal connectivity might be enhanced in VGPs. This conclusion was supported by looking at the regions that mostly changed their functional nodal strength between VGPs and NVGPs (**Fig 4D**). Thus, parieto-occipital and frontoparietal regions increased their functional nodal strength in VGPs the most. This local re-organization of brain connectivity suggests a possible increase in meso-scale integration [53] as consequence of expertise.

We also hypothesized that the functional re-organization observed in VGPs occurs in areas related to the cognitive demands of the RTS video games. We addressed this hypothesis using Neurosynth [49] and performed a term-based meta-analysis, allowing us to generalize our results from video games to other cognitive domains. Associations maps from diverse fMRI datasets [50] corresponded to the brain voxels that were consistently activated in studies associated with a specific term. We constrained our analysis to a list of 89 irreducible cognitive terms proposed by Poldrack et al. [51]. Then, we parcellated the maps using AAL90 and the resulting vectors, i.e., the regional associations obtained using Neurosynth were correlated with the difference in nodal strength between VGPs and NVGPs (the same vector used in **Fig 4D**). We show the top 10 correlations with cognitive terms in **Fig 4E**. After FDR correction, we found that the terms “reasoning” (*r* = 0.422, *p* = 0.004), “inference” (*r* = 0.374, *p* = 0.018) and “attention” (*r* = 0.362, *p* = 0.019) were positively correlated with the difference in nodal strength. Consequently, the brain areas that mostly increased their functional connectivity in VGPs are activated with these three cognitive domains. It is worth mentioning the positive correlation with the term “visual attention”, although it did not surpass the FDR correction.

Next, we investigated the functional subnetwork that correlates the most with playing time (**Fig 5**) using a greedy search algorithm. We included all the connections obtained using NBS (**Fig 4C**) as they represented the set of connections that are effectively different between VGPs and NVGPs. Then we correlated each entry of the masked FC with playing time, and the connection with the highest correlation was used as a base for adding more connections (**Fig 5A**). We repeated this procedure, adding more connections, until reaching a global maximum (**Fig 5A**). The subnetwork found with this algorithm has a mean FC strength that correlates positively with the playing time of VGPs (*r* = 0.570, *p* < 0.001, **Fig 5B**), and the network is also strengthened in VGPs with respect to NVGPs (*t* = 5.817, *p* < 0.001, **Fig 5B**). The functional subnetwork is presented in **Fig 5B**, and consists in a subset of parieto-occipital, frontal and motor brain areas, along with the posterior cingulate gyrus and the orbitofrontal section of the superior temporal pole. The ROIs are listed in **Table 2**.

**Fig 5.**
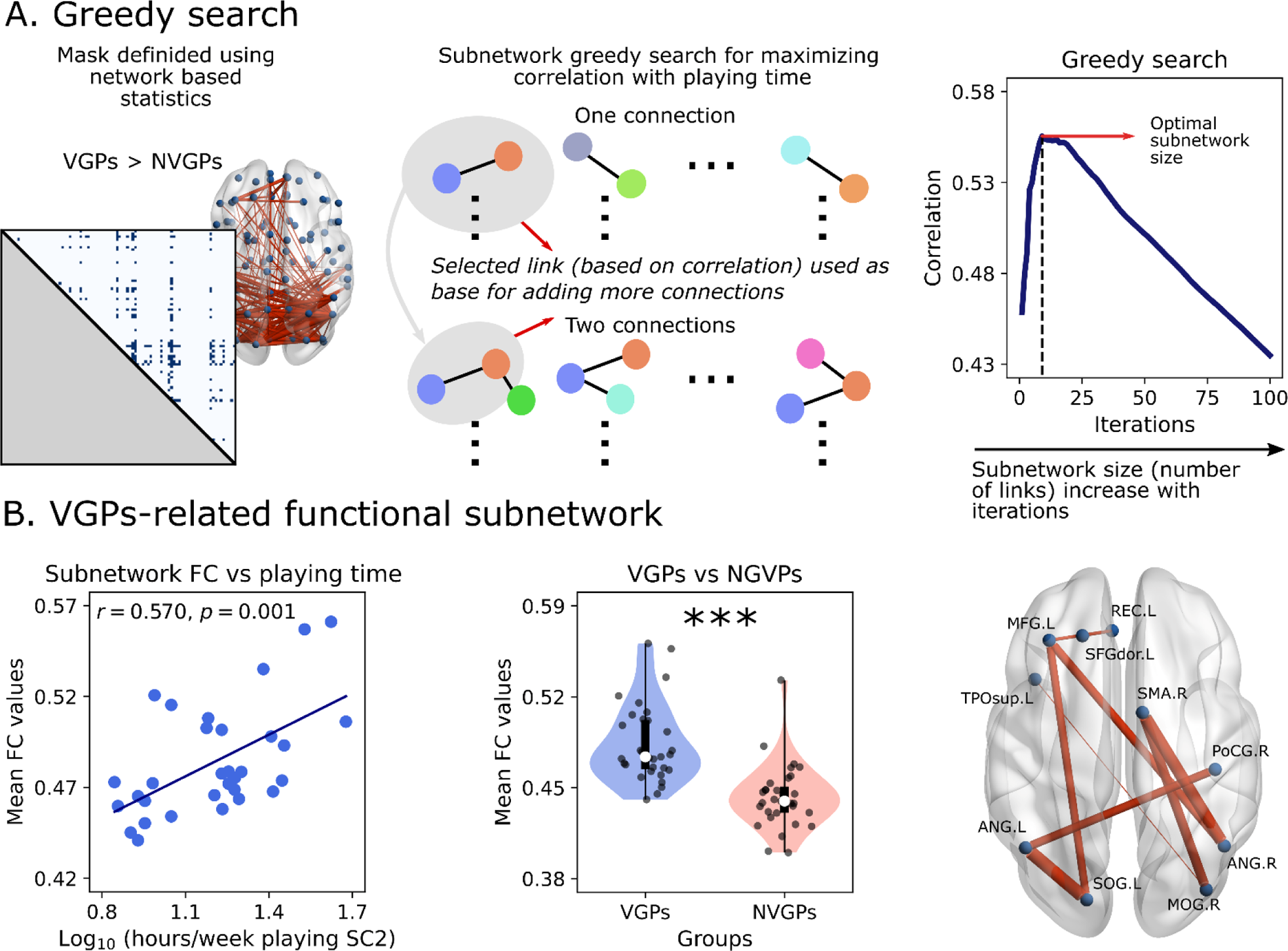
Greedy search for finding the minimal functional network associated with playing time. **A)** The greedy search algorithm was utilized with all functional connections identified by network-based statistics (NBS) as inputs. The algorithm first computed correlations between each functional connectivity (FC) value and playing time. The best connectivity value was chosen as the base for adding an additional connection, and this procedure was repeated until the global maximum was found. **B)** Correlation between the mean functional connectivity of the found subnetwork and playing time (left). This subnetwork also exhibited greater functional connectivity strength in video game players (VGPs) compared to non-video game players (NVGPs) (middle), enabling discrimination between the two groups. The functional subnetwork comprises occipital, parietal, frontal, and motor brain areas (right). Data points represent individual subjects (VGPs = 31, NVGPs = 31). Box plots were built using the 1st and 3rd quartiles, the median, the maximum and minimum values of distributions. The correlation was computed by Pearson’s correlation coefficient. ***: *p* < 0.001.

**Table 2.** List of ROIs obtained using the greedy search.

| Brain Regions |  |  |  |
| --- | --- | --- | --- |
| Left Hemisphere |  | Right Hemisphere |  |
| Name | Abbreviation | Name | Abbreviation |
| Angular gyrus | ANG.L | Angular gyrus | ANG.R |
| Middle frontal gyrus | MFG.L | Middle occipital gyrus | MOG.R |
| Rectus gyrus | REC.L | Postcentral gyrus | PoCG.R |
| Superior frontal gyrus (dorsolateral) | SFGdor.L | Supplementary motor area | SMA.R |
| Superior occipital gyrus | SOG.L |  |  |
| Temporal pole (superior) | TPOsup.L |  |  |

The ROIs are the same as the ones depicted in **Fig 5B**, and belong to the functional subnetwork that mostly correlates with playing time.

### 2.2 Gaming expertise increases robustness to noisy stimulation

We explored how the VGPs and NVGPs models respond to external stimulation [42–44]. Our aim was to test the models’ responsivity in a noisy context, that is, how much the noise influences the stimuli-mediated increase in connectivity and integration/segregation (**Fig 6**) to conceptualize the capability of the model of “filtering out” noise from stimuli. Specifically, we stimulated pairs of occipital homotopic brain areas, using noisy sinusoidal inputs (**Fig 6A**) and increasing the noise of the inputs progressively (**Fig 6A**). We measured the changes in FC and network topology, during stimulation, averaged across the six pairs of occipital ROIs.

**Fig 6.**
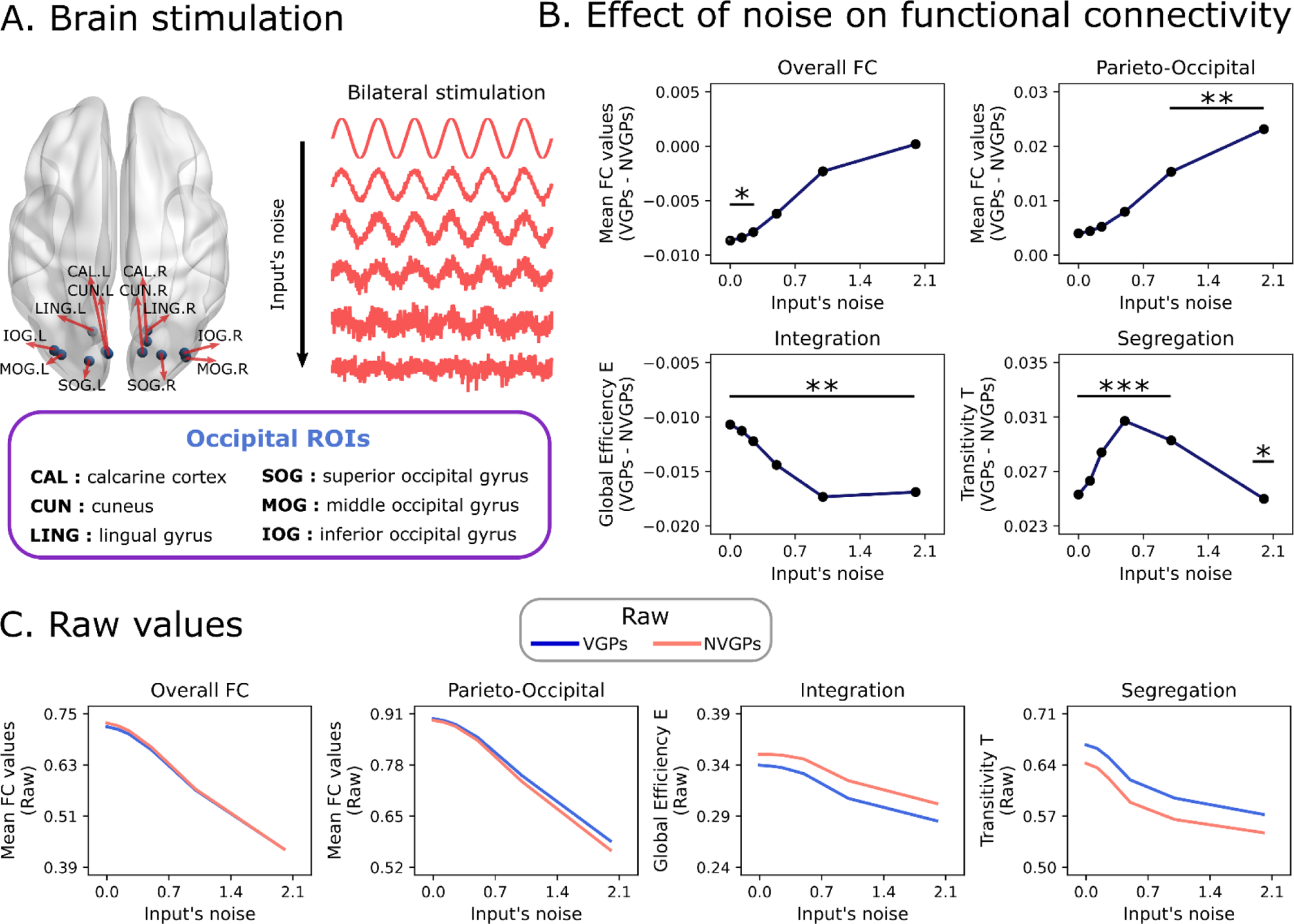
Responsivity of video game players and non-video game players to occipital stimulation. **A)** External stimulation was applied to pairs of homotopic occipital brain regions. The stimulation started with noise-free sinusoidal stimulation, and the level of input noise was increased from top to bottom. **B)** Difference on global functional connectivity (FC), parieto-occipital FC, integration (global efficiency) and segregation (transitivity) of video game (VGPs) and non-video game (NVGPs) players during stimulation. Lines with dots correspond to the difference between VGPs and NVGPs. **C)** Absolute values for VGPs and NVGPs. Statistical tests were performed using the single-subject models (VGPs = 31, NVGPs = 31). ***: *p* < 0.001, **: *p* < 0.01, *: *p* < 0.05. FDR corrected *p*-values were used.

While external stimulation always increased mean FC connectivity, increasing noise always reduced the global correlations, parieto-occipital FC, functional segregation, and integration (**Fig 6C**, where raw values for VGPs and NVGPs were plotted). However, VGPs and NVGPs responsivity to noisy inputs was not the same. VGPs parieto-occipital FC strength diverged from NVGPs when external stimulation becomes noisier. The specific differences comprised higher parieto-occipital FC in VGPs, in contrast to NVGPs, when input’s noise was subsequently increased in the model (**Fig 6B**). In the same way, the difference in functional integration and segregation becomes more pronounced when input’s noise is increased in the model (**Fig 6B**). The models’ perturbation indicates that occipital stimulation produces a shift to a more global segregated FC network topology, evidenced by the increase in functional segregation and parieto-occipital FC strength compared to RS conditions (compare with **Fig 4**). However, the meso-scale integration in VGPs is more robust in noisy contexts compared to NVGPs. In brief, our results suggest that video games expertise is both associated to functional reconfigurations in RS conditions, and with enhanced responsivity of the occipital ROIs to external stimulation.

### 2.3 Core structural connections characterize gaming expertise

The simulated FCs from VGPs and NVGPs only differ in the connectomes embedded into the model. Thus, FC differences emerge by the specific features of subjects’ connectomes. To provide a more direct link and to support previous results, we performed an additional in-silico experiment by converting NVGPs into VGPs (**Fig 7**). First, we fitted the whole-brain model using the averaged connectomes across subjects (averaged SSIM values of 0.427 for both VGPs and NVGPs). Then, we computed the difference matrix between VGPs and NVGPs SCs, and the connections were ranked from highest to lowest; top connections are the most different between VGPs and NVGPs. Next, we transferred a percentage of the ranked VGPs connections to the NVGPs’ averaged SC (**Fig 7A**). We compared the functional similarity between VGPs and “converted” VGPs (CVPGs) from the simulated FCs matrices (**Fig 7A**). These were first masked using the set of connections found using NBS (**Fig 4C** and **Fig 5A**). Then, the masked FCs were compared against each other (VGPs versus CVGPs) using the Euclidean distance. Lower distance values were expected when VGPs and CVGPs matrices started to converge to a similar FC pattern. Within each iteration of the algorithm, we increased the percentage of the structural connections transferred from the VGPs to CVGPs (**Fig 7A**). We found that the distance from VGPs decreased when progressively transferring connections to NVGPs (**Fig 7A**). We stopped when the distance from VGPs was stable as a function of the transferred connections (dashed lines in **Fig 7A**).

**Fig 7.**
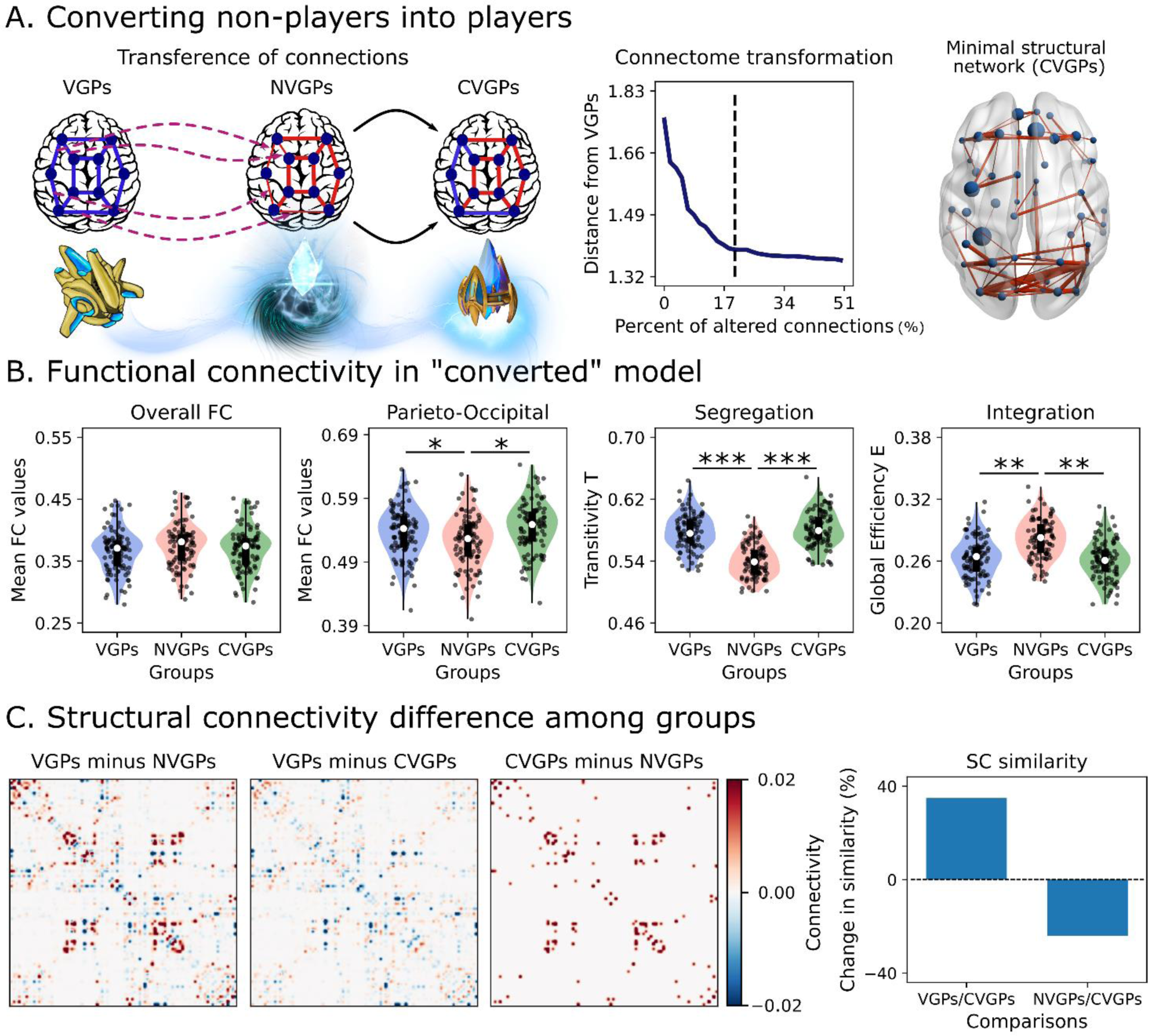
Converting non-players into players using the average model. **A)** The “conversion” procedure involved transferring the top structural connections (SC) from video game players (VGPs) to non-players (NVGPs). The connections that mostly differ between VGPs and NVGPs were transferred to NVGPs, creating a new group of converted VGPs (CVGPs). Then, we computed the distance from VGPs, as the Euclidean distance between the vectorized functional connectivity matrices of VGPs and NVGPs, previously reduced using the network-based statistics (NBS) mask. The minimal structural subnetwork associated with a stable reduction of the distance from VGPs is indicated by dashed lines in the plot and displayed in the rightmost panel. **B)** Global functional connectivity (FC), parieto-occipital FC, integration (global efficiency) and segregation (transitivity) of VGPs, NVGPs and CVGPs during resting-state conditions. **C)** Difference in structural connectivity among groups. The left panel shows the SC difference matrices, while the right panel illustrates the change in similarity of CVGPs in relation to VGPs and NVGPs’ SC matrices. Data points represent average model realizations (VGPs = 100, NVGPs = 100, CVGPs = 100 random seeds). Box plots were built using the 1st and 3rd quartiles, the median, the maximum and minimum values of distributions. ***: |D| > 1.2, **: |D| > 0.8, *: |D| > 0.5.

Only 20% of VGPs’ SC connections (considering SC entries greater than 0.01) had to be transferred to CVGPs for reproducing VGPs’ FC. The subnetwork found (**Fig 7A**) was the structural core of VGPs, that is, the minimal set of connections responsible for the differences between players and non-players. Interestingly, parieto-occipital and frontoparietal connections are present in the VGPs structural core, depicting connections between posterior brain areas with motor regions (i.e., supplementary motor area) and the anterior/posterior cingulate gyrus, and between the orbitofrontal cortex with parietal ROIs. The complete set of ROIs is listed in **Table 3**. The CVGPs’ functional connectivity properties resemble the ones described for VGPs (**Fig 7B**). We found negligible differences, between VGPs or CVGPs and NVGPs, in overall FC (D = −0.296 and D = −0.185 for comparisons with VGPs and CVGPs, respectively), moderate differences in parieto-occipital FC (D = 0.521, and D = 0.558), huge differences in global segregation (D = 1.723, and D = 1.891), and large differences in global integration (D = −1.015, and D = −1.162). Finally, we checked how SC changed in CVGPs when transferring the top 20% connections from VGPs. We show the difference between SC matrices in **Fig 7C**. Converting CVGPs increased the connectome similarity in relation to VGPs by 39.4%. In contrast, the conversion reduced the similarity to NVGPs by 24.1%. In consequence, the connectome conversion approaches CVGPs’ connectome to VGPs, but only partially. Despite that, CVGPs exhibited the full functional features of VGPs described before (**Fig 7B**). Thus, the structural core in players was critical and sufficient to reproduce the RS FC features of players.

**Table 3.** List of ROIs that comprise the minimal structural network of players.

| Brain Regions |  |  |  |
| --- | --- | --- | --- |
| Left Hemisphere |  | Right Hemisphere |  |
| Name | Abbreviation | Name | Abbreviation |
| Precentral gyrus | PreCG | Precentral gyrus | PreCG |
| Superior frontal gyrus (dorsolateral) | SFGdor | Superior frontal gyrus (dorsolateral) | SFGdor |
| Middle frontal gyrus | MFG | Superior frontal gyrus (orbital) | ORBsup |
| Inferior frontal gyrus (triangular) | IFGtriang | Middle frontal gyrus | MFG |
| Inferior frontal gyrus (orbital) | ORBinf | Inferior frontal gyrus (opercular) | IFGoperc |
| Supplementary motor area | SMA | Inferior frontal gyrus (triangular) | IFGtriang |
| Superior frontal gyrus (medial) | SFGmed | Supplementary motor area | SMA |
| Insula | INS | Superior frontal gyrus (medial) | SFGmed |
| Anterior cingulate gyrus | ACG | Rectus gyrus | REC |
| Median cingulate gyrus | MCG | Anterior cingulate gyrus | ACG |
| Parahippocampal gyrus | PHG | Median cingulate gyrus | MCG |
| Calcarine cortex | CAL | Calcarine cortex | CAL |
| Cuneus | CUN | Cuneus | CUN |
| Lingual gyrus | LING | Lingual gyrus | LING |
| Superior occipital gyrus | SOG | Superior occipital gyrus | SOG |
| Middle occipital gyrus | MOG | Middle occipital gyrus | MOG |
| Fusiform gyrus | FFG | Inferior occipital gyrus | IOG |
| Postcentral gyrus | PoCG | Fusiform gyrus | FFG |
| Superior parietal gyrus | SPG | Postcentral gyrus | PoCG |
| Inferior parietal gyrus | IPG | Superior parietal gyrus | SPG |
| Angular gyrus | ANG | Inferior parietal gyrus | IPG |
| Precuneus | PCUN | Supramarginal gyrus | SMG |
| Paracentral lobule | PCL | Angular gyrus | ANG |
| Putamen | PUT | Precuneus | PCUN |
| Superior temporal gyrus | STG | Caudate | CAU |
| Temporal pole (superior) | TPOsup | Superior temporal gyrus | STG |
| Middle temporal gyrus | MTG | Temporal pole (superior) | TPOsup |
| Middle temporal gyrus | MTG |  |  |
| Temporal pole (middle) | TPOmid |  |  |
| Inferior temporal gyrus | ITG |  |  |

Finally, we repeated the same graph theoretical analyses using the SC instead of FC (**Fig 8**). We observed that neither whole-brain number of fibers (*t* = 1.038, *p* = 0.404), parieto-occipital number of fibers (*t* = 2.127, *p* = 0.150), structural integration (*t* = 1.490, *p* = 0.283), and segregation (*t* = 0.478, *p* = 0.634), were different between VGPs and NVGPs. The NBS analysis suggested no differences in VGPs and NVGPs brain networks (*p* = 0.290 using 10000 permutations). However, with no FDR-correction, the number of fibers within the parieto-occipital network was higher in VGPs than in NVGPs with statistical significance (*t* = 2.127, *p* = 0.038, which correspond to the original report in Kowalczyk-Grębska et al. [9]). These results confirm the importance of looking at the FC to characterize the differences associated with expertise, supporting the approach used in our work.

**Fig 8.**
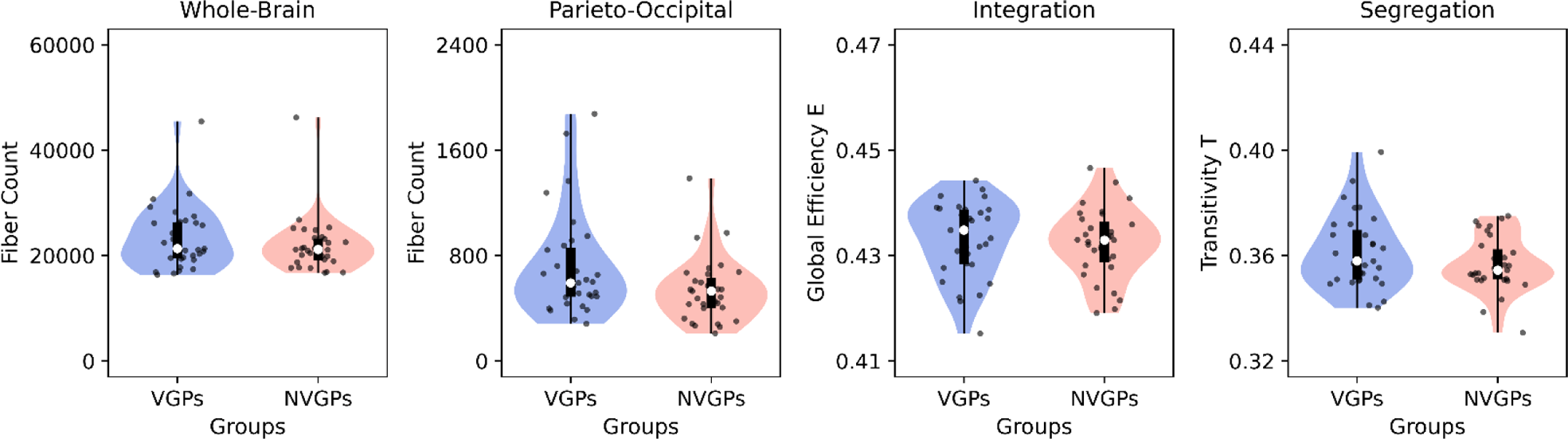
Structural connectivity of players and non-players. **A)** Number of fibers at the whole-brain level and within the parieto-occipital subnetwork, integration (global efficiency) and segregation (transitivity) of video game (VGPs) and non-video game (NVGPs) players. Data points represent individual subjects (VGPs = 31, NVGPs = 31). Box plots were built using the 1st and 3rd quartiles, the median, the maximum and minimum values of distributions. ***: *p* < 0.001, **: *p* < 0.01, *: *p* < 0.05. FDR corrected *p*-values were used.

## 3 Discussion

We used a whole-brain semi-empirical model to unveil the structural neural plasticity mechanisms associated with gaming expertise. We inferred VGPs and NVGPs FC using whole-brain simulations (data completion) and predict brain responsivity of VGPs under noisy contexts by in-silico simulation (uncovering latent information). This approach allowed us to integrate multimodal neuroimaging data into single subjects’ models, characterize brain dynamics both in RS and during stimulation, and generalize the findings from video games expertise to other cognitive domains using Neurosynth [31,54]. Findings suggest that gaming expertise induced neural plasticity at structural connectivity that in turns induces functional meso-scale re-organization in the brain. Repeating the graph theoretical analyses at the level of SC showed no differences between VGPs and NVGPs, remarking the usefulness of our approach looking at FC for characterizing brain connectivity differences associated with expertise. The results shed light about how video expertise can reshape FC through structural neural plasticity mechanisms, and how these alterations are related to cognitive functions.

A shift to a more locally integrated network re-organization was observed in VGPs. This meso-scale integration [30] consisted in a decrease in global efficiency (integration) and in an increase in global clustering coefficient (segregation). Notably, these are differences in the global properties of the brain network topology, that were not found in the previous connectome analysis [37] and emerge in FC from the model informed with SC. Consequently, connectivity strength was enhanced in specific subnetworks. Increased network segregation has been reported in different behavioral paradigms, including visual attention [55], motor learning [56] and sustained attention [57] tasks. Local connectivity was found in salience, cingulo-opercular, dorsal attention and visual networks. The functional re-organization towards meso-scale integration is also observed by neuromodulation [30,58], in terms of context-dependent effect [59] of cholinergic and noradrenergic systems on FC. In contrast with these transient changes on network topology produce by neuromodulatory systems, the structural “re-wiring” associated with video game training might be long-standing (not transient), possibly increasing the baseline bandwidth of communication of frontoparietal and parieto-occipital networks. Thus, our results and previous reports are consistent with the idea that a more global segregated network topology might prevent the cross-interference between brain networks, leading to a more precise processing of task-relevant information within the brain [60].

Expertise can lead to mechanistic changes in white and grey matter structures directly required for specific tasks. At the micro-structural level, long term potentiation has been proposed as a mechanism involved in learning-based neural plasticity [61], reenforcing synaptic strength of neurons co-activated together. However, recent studies suggest that white matter neural plasticity plays a pivotal role in enhancing communication of brain regions involved in learning, experiences, and expertise [62,63]. Activity-dependent axonal propagation can lead to white matter structural changes driven by oligodendrocytes, e.g., modulation of myelin thickness, that can alter action potential transmission speed across axonal fibers [62,63]. Although our model does not include transmission delays explicitly, similar computational models showed that both structural connectivity strength and transmission delays alters the likelihood of synchronization between brain areas [64]. Thus, in our model a higher density of axonal fibers can be interpreted as increased communicability or connectivity strength between brain regions. In the context of RTS video games, brain regions involved in visual attention, logic inference, memory, motor outputs and reasoning should coordinate their activity for optimal in-game performance. Within that context, the increased connectivity strength of frontoparietal and parieto-occipital brain regions in VGPs –meso-scale integration– would allow a more effective transfer of information between and within these subnetworks. Indeed, these subnetworks have been consistently associated with enhanced performance in attentional and working memory tasks [65], and were shown to be highly plastic [66,67], more specifically, were shown to be susceptible to functional and structural (white matter) reshaping as consequence of training and stimulation.

RTS video games are unique in the way that they not only require the use of logical reasoning and memory to anticipate other players’ actions, but also involve real-time coordination and decision-making, demanding quick and accurate choices. The fMRI meta-analysis using Neurosynth revealed positive correlations between regional functional strength (VGPs minus NVGPs) with the association’s maps of attention, reasoning, and inference cognitive terms. That is, brain areas with increased functional strength in VGPs are also consistently activated in fMRI studies related to these concepts. The regions that we found using the greedy search and connectome transformation are suggested to be involved in these specific cognitive skills [1,5,6,65]: 1) identified by greedy search: angular gyrus, posterior cingulate gyrus, supplemental motor area, and frontal and parieto-occipital brain regions; 2) identified by the connectome transformation: the supplemental motor area, precuneus, cuneus, anterior and posterior cingulate gyrus, hippocampus and parahippocampal gyrus, with other frontal and parieto-occipital areas. These frontoparietal and parieto-occipital regions are essential to RTS games performance [1,5–8,11,12,68], and our results suggest that structural neural plasticity might be the mechanism responsible for the increased connectivity between them. Previous work in the field also supports our findings. For example, strategy video games are associated with hippocampal and prefrontal cortex structural changes [69], and enhanced FC between frontal and visual regions [8]. On the other hand, action video games are linked to increased functional integration between attentional and sensorimotor networks [70], necessary for visual attention and muscle coordination [65].

We tested the VGPs response to stimulation, to mimic the occipital activations in response to visual stimuli. We found that stimulation of occipital ROIs produced a higher increment of parieto-occipital FC in VGPs, compared to NGVPs, when inputs become noisier. Similar results hold for integration and segregation. As previously reported [9], VGPs connectomes are characterized by higher parieto-occipital SC strength. We suggest that neural plasticity increased connectivity strength within the parieto-occipital brain network, as consequence of RTS video games expertise, and this decreased the influence of noise in information processing [17]. Certainly, information transfer gets easier within densely interconnected brain networks; modulation and stimulation of brain regions is more effective in coordinating brain activity when nodes belong to the top strength brain areas [71,72]. In that way, efficient coding theory proposed that visual system exploits information redundancy to increase signal-to-noise ratio of visual stimuli [20,21]. An architecture of dense connections within the parieto-occipital loop may support efficient coding within visual cortex [33]. Thus, our results suggest that VGPs’ SC not only strengthened parieto-occipital FC in RS conditions, but also enhances FC more efficiently in the context of brain stimulation. It may be possible that the bandwidth of communication within the parieto-occipital loop is higher in VGPs than NVGPs, and/or VGPs SC increases signal-to-noise ratio of the incoming visual stimuli. The speed of processing of these stimuli is closely related to FC between frontoparietal and visual networks [73], and it has been reported that video game players have improved decision-making and response times than non-players [1,68]. Enhanced communication of parieto-occipital and frontoparietal brain networks might be essential for making efficient and faster decision in RTS games, where the reaction time is crucial to adapt, as fast as possible, one’s own strategies in response to other players’ actions.

Our work emphasizes the role of computational models in reinforcing the findings of empirical studies. By inferring VGPs and NVGPs FC data from whole-brain modeling [24,26], we linked structural neural plasticity to the functional alterations in VGPs, despite subjects’ functional data being not available. Further, from RS conditions, latent information could be uncovered using models [24], as was done in this work with in-silico stimulation, one of the novelties that differentiates our work from other approaches in the field. Moreover, our work suggests that VGPs have stronger FC strength in networks affected in some brain disorders. For example, in mild cognitive impairment and Alzheimer’s disease consistent pathophysiological disturbances are increased atrophy and loss of gray matter volume in parieto-occipital and frontoparietal brain networks [74]. As suggested by our results, expertise in RTS games might produce plastic changes in these subnetworks, enhancing their FC strength. In that way, video game training might be used as a complementary therapy in neurodegenerative diseases [1,3], with the aim to restore the healthy spatial-temporal dynamics of patients. In that line, video game training has demonstrated a promising therapeutic potential in amblyopia [1]. As video games are associated with improved attention and cognition [1,5,6], they might be used for dampening the cognitive decline associated by aging [4,75]. Finally, video games constitute a more engaging, motivating, rewarding, attractive and naturalistic tool that can be used to complement clinical interventions, fundamental to ensure the continuity of patients’ interventions.

We must point out some limitations of our results. First, we are aware of the small sample size used in this work. Further validation of our results should be confirmed using larger data sets. However, we take advantage of whole-brain modeling by performing several realizations (simulations) for each individual subject, reducing measurements variability, and ultimately making a better use of small datasets [76]. Even though the lack of functional data (e.g., EEG, fMRI) is a weakness, we used a whole-brain model to test the predictions about FC of players. Another limitation of the data is missed in-game scores, for example, the number of actions per minute of VGPs, that might constitute better descriptor of players’ in-game performance than the weekly playing time. Further, our cross-sectional study cannot allow us to make a direct causal connection between video games expertise with differences in FC [1]. Finally, further work would incorporate explicit neural plasticity mechanisms in the model [77], to directly test how brain stimulation shapes anatomical connections.

Together, our results suggest that expertise-driven structural neural plasticity may be responsible for the functional meso-scale integration associated with RTS games expertise, leading to increased functional connectivity in frontoparietal and parieto-occipital brain regions. This network re-configuration would be related to more efficient early and late processing of stimuli. Our work constitutes a novel contribution to understanding how neural plasticity mechanisms are associated with meso-scale functional connectivity, and how these changes can be linked to general cognitive skills acquired with expertise.

## 4 Material and Methods

### 4.1 Participants and ethics statement

Sixty-four right-handed male subjects participated in this study. Two subjects were excluded from the analysis because of the bad quality of their MRI data, so the final sample was 62 participants. All subjects completed an online questionnaire about demographics, education level and video game playing experience (questionnaire description – detailed description in [9,78]). Inclusion criteria for the RTS group, *n* = 31 (mean age = 24.7 years, SD = 4.27), were as follows: (a) experienced in RTS and StarCraft II game playing, (b) played RTS games at least 6 hours/week for the past 6 months, (c) declared playing StarCraft II for more than 60% of total game play time, and (d) was an active player (played matches in last two seasons) and was placed in one of five StarCraft Leagues (Gold, Platinum, Diamond, Master, Grandmaster). The inclusion criteria for NVGPs, *n* = 31 (mean age = 24.4 years, SD = 3.00), were as follows: (a) less than 6 hr of RTS video game play, and (b) less than 8 hours a week of total video game play (across all genres, including RTS) over the past 6 months. Only male participants were recruited due to the lack of female participants with adequate video game experience. See **Table 4** for complete demographic information.

**Table 4.**
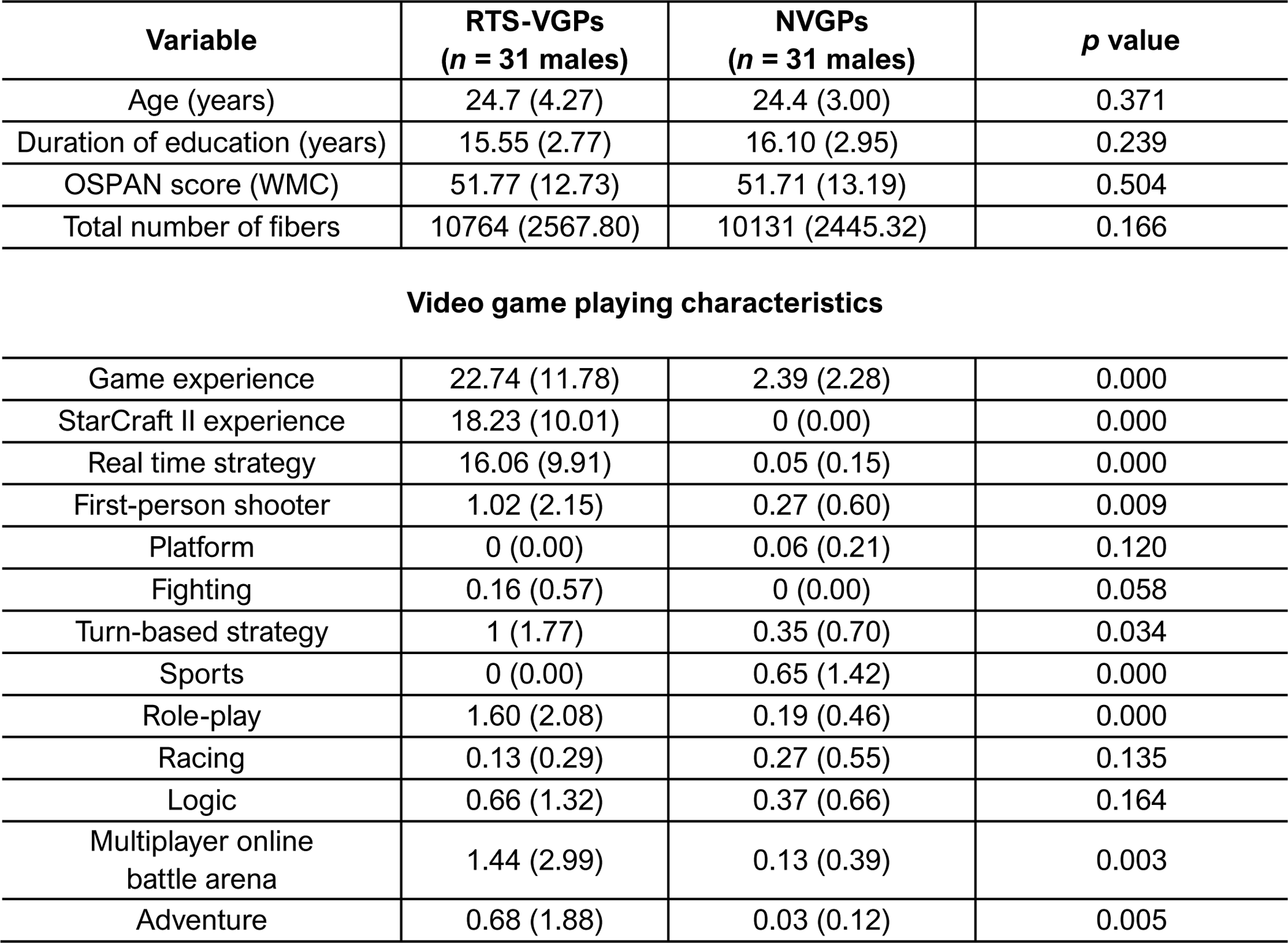
Participants characteristics. Demographic and white matter properties of all participants.

| Variable | RTS-VGPs<br>(*n* = 31 males) | NVGPs<br>(*n* = 31 males) | *p* value |
| --- | --- | --- | --- |
| Age (years) | 24.7 (4.27) | 24.4 (3.00) | 0.371 |
| Duration of education (years) | 15.55 (2.77) | 16.10 (2.95) | 0.239 |
| OSPAN score (WMC) | 51.77 (12.73) | 51.71 (13.19) | 0.504 |
| Total number of fibers | 10764 (2567.80) | 10131 (2445.32) | 0.166 |
| **Video game playing characteristics** |  |  |  |
| Game experience | 22.74 (11.78) | 2.39 (2.28) | 0.000 |
| StarCraft II experience | 18.23 (10.01) | 0 (0.00) | 0.000 |
| Real time strategy | 16.06 (9.91) | 0.05 (0.15) | 0.000 |
| First-person shooter | 1.02 (2.15) | 0.27 (0.60) | 0.009 |
| Platform | 0 (0.00) | 0.06 (0.21) | 0.120 |
| Fighting | 0.16 (0.57) | 0 (0.00) | 0.058 |
| Turn-based strategy | 1 (1.77) | 0.35 (0.70) | 0.034 |
| Sports | 0 (0.00) | 0.65 (1.42) | 0.000 |
| Role-play | 1.60 (2.08) | 0.19 (0.46) | 0.000 |
| Racing | 0.13 (0.29) | 0.27 (0.55) | 0.135 |
| Logic | 0.66 (1.32) | 0.37 (0.66) | 0.164 |
| Multiplayer online<br>battle arena | 1.44 (2.99) | 0.13 (0.39) | 0.003 |
| Adventure | 0.68 (1.88) | 0.03 (0.12) | 0.005 |

The NVGP sample was characterized by no experience with StarCraft II, and very little video game experience across genres. Based on a preliminary self-report questionnaire, StarCraft II gaming experience was verified by the activity from the last two seasons (number of games played). The groups were matched in terms of education level as all participants’ education were undergraduate level. What is more, we controlled participants’ working memory capacity (WMC) using the operation span task (OSPAN [79], and detailed description in [9]). None of the participants had a history of neurological illness and they did not use psychoactive substances. All subjects provided a written informed consent to participate in the experiment, and the study protocol was approved by the SWPS University Ethical Committee in accordance with the Declaration of Helsinki. All subjects participated in additional MRI sessions and cognitive measurements, which were not related to the project described in this article.

Mean and standard deviation (SD in parentheses) represent the values. A nonparametric permutation test was employed to compare all variables between the groups. The term ‘Game experience’ refers to the average number of hours per week spent playing video games in the last 6 months. Similarly, “StarCraft II experience” indicates the average number of weekly hours spent specifically on playing the StarCraft II game over the past 6 months (for further details, refer to the Material and Methods section). Table adapted from Kowalczyk-Grębska et al. [9].

### 4.2 MRI acquisition and preprocessing

MR images were collected on a 3-Tesla MRI scanner (Siemens Magnetom Trio TIM, Erlangen, Germany) equipped with a 32-channel phased array head coil. All subjects were also informed to minimize their head movements during the scanning procedure. First, anatomical data of the brain were acquired from T1-weighted images using an MPRAGE sequence with the following parameters: repetition time [TR] = 2530 ms; echo time [TE] = 3.32 ms; flip angle = 7°; 176 slices; voxel size = 1 x 1 x 1 mm^3^. Next, a Spin-echo diffusion weighted echo planar imaging (DW_EPI) sequence was performed with TR = 8700 ms; TE = 92 ms; GRAPPA = 2; flip angle = 90°, voxel size = 2 x 2 x 2 mm^3^, 64 gradient directions with b-value of 1000 s/mm, along with two images with no diffusion gradient applied (b-value = 0). The DWI sequence was repeated to increase signal to-noise ratio.

### 4.3 DTI Data preprocessing and structural networks

DTI image preprocessing was implemented using the “Pipeline for Analyzing Brain Diffusion Images’’ (PANDA) software [80]. DICOM files (64 directions) were first converted into a single four-dimensional NIFTI format. All diffusion-weighted images were visually inspected for artifacts due to subject head motion. Next, eddy current induced distortion as well as simple head-motion artifacts were corrected by applying affine alignments of the diffusion-weighted images to the average of b0 images [81]. Diffusion tensors were constructed using weighted-least-squares. Following, diffusion tensors as well as FA matrices were calculated for each participant using the DTIFIT tool [82,83]. Using the affine transformation, the individual FA image in native space was co-registered to its corresponding structural image (T1-weighted). Each individual structural image was then nonlinearly registered to the MNI space using the MNI152_T1_2mm_brain template, and the resulting transformation was inverted and used to warp the AAL atlas [84] from the MNI space to the T1 space. The individual structural image (T1-weighted) was co-registered to its corresponding FA image in native space using the affine transformation, and the AAL atlas was thus warped to the individual FA space. We employed the AAL atlas here to parcellate the individual cerebrum into 90 brain regions (45 for each hemisphere; regions listed in **Table 1**) defined based on local gyri and sulci patterns. The whole-brain streamline deterministic fiber tractography was then performed for each image by using the diffusion tensors to compute the fiber tracts [85,86]. The maximum turning angle was 45 degrees and FA threshold was 0.2. The resulting white matter fiber tracts for whole brain were reviewed in Trackvis software (http://trackvis.org/dtk/). Network matrix was constructed by calculating the number of fibers connecting each pair of brain regions [87–89] for all subjects, where the brain regions serve as nodes and fiber numbers serve as edges. The resultant 90 x 90 symmetric matrices, one for each participant, were finally used in network analysis. The matrices were normalized between 0 and 1, dividing each individual matrix by its global maxima.

### 4.4 Empirical functional data

The functional neuroimaging dataset utilized in this study was obtained from the Human Connectome Project (HCP, http://www.humanconnectome.org/), Release Q3. As per the HCP protocol, all participants provided written informed consent to the HCP Consortium. This dataset included fMRI acquisitions from 100 unrelated subjects, who were part of the HCP 900 data release [39]. All HCP scanning procedures were authorized by the Institutional Review Board at Washington University in St. Louis.

HCP dataset employed consisted in a 3D MPRAGE T1-weighted structural MRI with TR of 2400 ms, TE of 2.14 ms, TI of 1000 ms, flip angle of 8°, FOV of 224 × 224, and voxel size of 0.7 mm isotropic. Additionally, two 15-minute resting-state fMRI sessions were performed using gradient-echo EPI with TR of 720 ms, TE of 33.1 ms, flip angle of 52°, FOV of 208 × 180, and voxel size of 2 mm isotropic. Only the first scanning session in the LR direction was used for functional data. We used the HCP ICA-FIX denoised fMRI RS preprocessed data. The preprocessing pipeline consisted in several steps, including distortion correction to remove spatial distortions, volume realignment to account for subject motion, registration of the fMRI data to the structural image, bias field reduction, normalization of the 4D image to a global mean, and masking of the data using the final brain mask. The pipeline also incorporates ICA-FIX for denoising [90–92]. The resulting output of this pipeline can be used for standard volume-based analyses of the fMRI data. The fMRI RS volumetric time series were parcellated using the AAL90 brain parcellation [40], which considered *n* = 90 cortical and subcortical brain regions. After parcellation, they were band-pass filtered between 0.01 and 0.1 Hz with a Bessel third-order filter [93,94].

### 4.5 Whole-brain modeling and simulations

Our whole-brain model combines single-subject empirical SC, HCP empirical FC, and regional dynamics that rules the behavior at the single region level. We considered a network of *n* = 90 brain areas defined by the AAL90 parcellation (**Fig 2A**). For simulating the local brain activity, we used the normal form of a supercritical Hopf bifurcation (Stuart-Landau oscillators) [28,29]. This bifurcation can alter the nature of the system’s trajectories from a limit cycle, that yields self-sustained oscillations, to a stable fixed point in phase space (**Fig 2B**). The whole-brain computational model comprises a set of model parameters that govern the global dynamical behavior. Among these, the global coupling, *G*, represents the global conductivity of the fibers scaling the SC between brain areas, which is assumed to be equal across the brain [28,29,95]. The other crucial parameters are nodal local bifurcation parameter, *a*_*i*_, which governs the dynamical behavior of each area *i* between noise-induced oscillations (*a*_*i*_ < 0), self-sustained oscillations (*a*_*i*_ > 0), or a critical behavior between both (*a*_*i*_ ∼ 0) (**Fig 2B**). The brain areas also receive uncorrelated gaussian noise, *η*_*i*_(*t*), with *β* = 0.1 standard deviation. The local dynamics of each brain area was described by the following set of ordinary differential equations

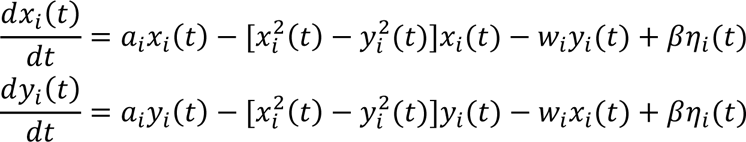

In the equations, *x*(*t*) emulates the BOLD-like signals (real component of time series), while *y*(*t*) corresponds to the imaginary component. The regions presented self-sustained oscillations at a frequency *f*_*i*_ = *w*_*i*_/2*π* for *a*_*i*_ > 0, and noise-driven oscillations for *a*_*i*_ < 0. We set the frequency of oscillation *f*_*i*_ = 0.05 Hz for overall nodes. Brain areas were also coupled using empirical structural connectivity matrices (weighted and undirected), denoted by *M*. Each entry *M*_*ij*_ indicates the existence of a connection between regions *i* and *j* and its strength. The entries of *M* scales with the global coupling, *G*, leading to the next set of equations

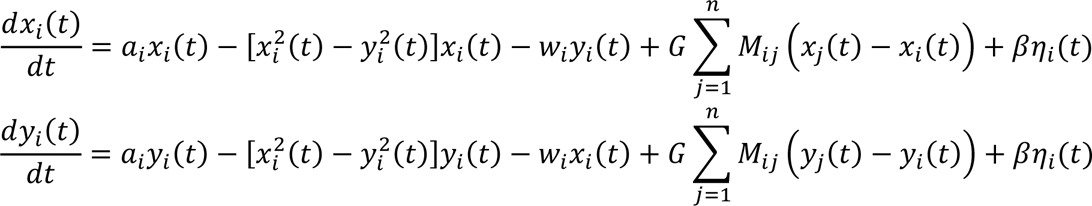

Further, the model incorporates external perturbation (stimulation, **Fig 2B**), following previous works using the Hopf model [42,43]. The perturbation consisted in an external additive periodical forcing term in the equations

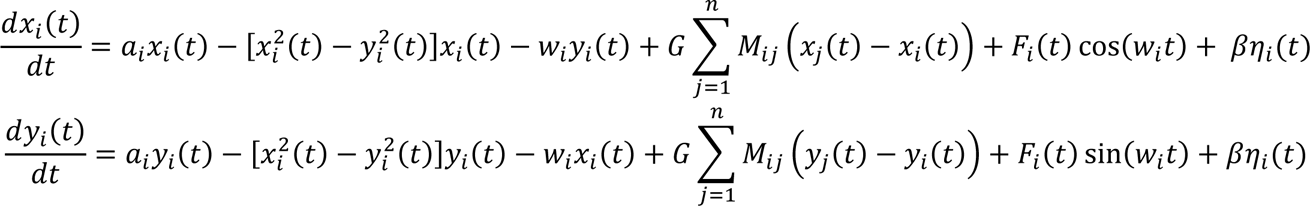

with amplitude *F*_*i*_(*t*) = *F*_0*i*_ + *σμ*_*i*_(*t*), with *F*_0*i*_ the amplitude of the sinusoidal perturbation, and *μ*_*i*_(*t*) uncorrelated gaussian noise with *σ* standard deviation. For the RS condition, model fitting and connectome transformation (see subsections below), we set *F*_0*i*_ = 0 and *σ*_*i*_ = 0 across nodes.

We performed 15 min simulations using an integration step of 100 msec, using the Euler-Maruyama integration scheme for stochastic differential equations. The BOLD-like time series were band-pass filtered between 0.01 and 0.1 Hz, as the empirical fMRI BOLD RS data from other works in the field [93,94].

### 4.6 Model fitting

#### 4.6.1 Single-subject model

First, we computed the empirical FC, which consisted in the average FC across the 100 HCP subjects and RS periods. This FC matrix constituted the objective function for model fitting. We informed our model with subject-specific SC matrices of VGPs and NVGPs. For each single-subject model, we swept the global coupling parameter, *G*, between 0 and 3 in increments of 0.05. The simulated BOLD-like time series were used for building the simulated FC matrices (**Fig 3A**). Then, empirical and simulated FCs were compared against each other using the Structural Similarity Index (SSIM) [45] (**Fig 3A**). The SSIM measures the similarity between two images or video frames, for example, between two FC matrices. It is a full-reference image quality assessment method that considers the perceived changes in structural information, luminance, and contrast between the reference and distorted images. The SSIM has been used in other whole-brain modeling works [44,46]. It is bounded between 0 (poor fitting) and 1 (good fitting). For each subject and *G* value, we used 100 random seeds, reporting the average SSIM across seeds. Finally, we found a single optimal *G* value, which maximizes the SSIM, for each one of the single-subject models (**Fig 3B**). These *G* values were fixed across future simulations.

#### 4.6.2 Averaged model

We also fitted an average version of the model, using the mean SC matrices for VGPs and NVGPs. The averaged VGPs and NVGPs models were used for connectome transformation (see the corresponding subsection below). The procedure was similar to the single-subject fitting, but instead we used the averaged SCs and unique values of *G* were found for VGPs and NVGPs.

### 4.7 Functional connectivity network analysis

First, we built the FC matrices using pairwise Pearson’s correlation between regional time series. From the FCs, we computed the global correlations (overall FC) as the mean values within the FC matrices. Also, we calculated the average correlations of parieto-occipital subnetwork, using the ROIs’ labels of the AAL90 parcellation as a mask.

For the analysis of topological properties of functional networks, we applied a proportional thresholding to FCs for removing any spurious connectivity values, as outlined by van den Heuvel et al. [96]. Afterward, the matrices were binarized, and graph metrics were computed. Specifically, we calculated the binary global efficiency and transitivity metrics, which are related to integration and segregation, respectively, according to Rubinov & Sporns [47].

Global efficiency, which is based on paths, was defined as [97]:

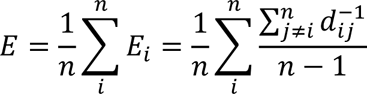

where *E*_*i*_ represents nodal efficiency, *n* = 90 the total number of nodes, *d*_*ij*_ the shortest path that connects nodes *ij* and *E* is the average nodal efficiency across nodes (network global efficiency). The metric ranges between 0 and 1, with higher values indicating greater network integration, where distant nodes can communicate easily, and values near 0 indicating the opposite. Transitivity, on the other hand, is based on the count of triangular motifs in the network and was computed using the formula proposed by Newman [48]:

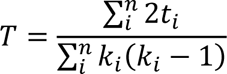

with *t*_*i*_ as the number of triangles around the node *i*, and *k*_*i*_ as the node degree. The number of triangles is defined as:

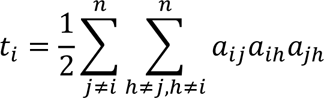

where *a*_*ij*_ = 1 if nodes *ij* are connected in the graph, and 0 otherwise. Transitivity measures the extent of local (short-range) interactions between nodes and, in simpler terms, reflects how much nodes shared common neighbors between them. Higher transitivity values are expected in segregated networks, whereas values close to 0 are expected in completely random networks. Transitivity is a variant of the clustering coefficient, differing only in its normalization procedure: while transitivity is normalized collectively, the clustering coefficient is normalized at the single node level, as described by Rubinov & Sporns [47].

To avoid the potential bias of using a single proportional threshold, we applied a range of thresholds from 5% to 18% with linear increments of 1%. The area under the curve (AUC) of each graph metric was reported for the explored threshold range, as suggested by Ginestet et al. [98]. The threshold range was established based on exploratory analyses, as a stable difference (VGPs - NVGPs) was found across network densities. Also, we computed global correlations (overall FC), for verifying that the differences in functional network topology were not ascribed to overall FC [96]. We performed the graph analysis using the Brain Connectivity Toolbox for Python (https://github.com/fiuneuro/brainconn) [47].

In addition to the global network analysis outlined in the previous sections, we utilized network-based statistics (NBS) to identify key connections that are different between VGPs and NVGPs [52]. This method detects subsets of links that exhibit differences between two conditions and circumvents the issue of multiple comparisons that arises when conducting paired tests on individual links. We used the NBS implementation of the Brain Connectivity Toolbox for Python (https://github.com/fiuneuro/brainconn) [47,52] and conducted a one-tailed paired *t*-test to rank the connections, for identifying the strengthened connections in VGPs. We chose a threshold of *t* = 3 to capture the most significantly altered connections caused by video game expertise. Subsequently, we determined the size of the largest component, and evaluated the statistical significance of the identified subnetworks using permutation tests, which entailed calculating the size of the largest component for each permutation (up to 10000 permutations).

We repeated the same graph theoretical analyses for SC but reducing the threshold for NBS from of *t* = 3 to of *t* = 1.5. The rest remained unchanged.

### 4.8 Neurosynth associations

Neurosynth is a neuroscience tool that uses automated meta-analysis to identify the most frequently activated brain regions associated with specific concepts or cognitive processes. It integrates large-scale neuroimaging data from thousands of studies to generate probabilistic maps of brain activation patterns associated with different mental states, behaviors, and stimuli [49]. We obtained the associations maps ascribed to a predefined list of 89 irreducible cognitive terms proposed by Poldrack et al. [51]. Then, the maps were parcellated using AAL90. We used Neurosynth to find correlations between the mean difference in nodal strength (VGPs minus NVGPs) and the association maps into the AAL90 parcellation.

### 4.9 Greedy search

We performed a greedy search to find the FC subnetwork that mostly correlates with playing time (hours per week). We considered a mask of connections, previously reported using NBS (**Fig 5A**). That subnetwork found using NBS consisted in the set of connections that mostly increase their strength in VGPs compared to NVGPs. Then, Pearson’s correlation between all connections and playing time was computed, and the connection associated with the highest correlation was used as base for adding more connections (**Fig 5A**). More connections were added one by one, until correlation with playing time reached a global maximum.

### 4.10 Perturbations

We analyzed the VGPs and NVGPs models robustness to noisy external stimulation. Using the single-subject models, we fixed the magnitude of external stimulation to *F*_0*i*_ = 0.5, and several input’s noise level, *σ*_*i*_, were considered from 0 to 2 (**Fig 6A**). Following previous works in the field [42,43], we stimulated pairs of homotopic brain regions, constraining our simulations to occipital ROIs (**Fig 6A**). We compared how the overall FC, parieto-occipital FC, integration, and segregation differences between VGPs and NVGPs during stimulation by different noise levels.

### 4.11 Connectome transformation

For finding a direct link between the functional differences of VGPs in relation to NVGPs, we performed a “connection transformation” starting from the NVGPs averaged model. We first ranked the stronger connections of VGPs in contrast to NVGPs. Then, a percentage of the top VGPs’ ranked connections were added to the NVGPs averaged connectome, creating the converted VGPs (CVGPs) group (**Fig 7A**). After transferring connections, CVGPs mean SC was normalized respect to NVGPs, to preserve the averaged SC strength between groups. We assessed the functional similarity between VGPs and CVGPs computing the Euclidean distance between the NBS-based FC (**Fig 5A**) of both groups. The set of connections associated with a stable reduction of the Euclidean distance constitutes the minimal subnetwork responsible for FC in players.

### 4.12 Statistical analysis

Groups comparisons for demographic data, white matter tractography and video-game characteristics were reported as in the original publication of Kowalczyk-Grębska et al. [9], where nonparametric permutation tests implemented in the GRETNA package were used to assess groups differences [99]. Permutation tests are robust for small sample sizes and are well-suited when normality assumption is not guaranteed [100]. The test consisted of computing the mean difference between groups, and then computed the computed value with a distribution of mean differences obtained by a random shuffling of groups’ labels, up to 10000 permutations. From the single subject whole-brain models, statistical comparison between VGPs and NVGPs consisted in pairwise *t*-Student tests from independent samples, assuming a critical *p*-value of 0.05 for rejecting the null hypotheses. All the statistical relationships were assessed using Pearson’s correlation. The resulting *p*-values of multiple comparisons and correlations were FDR corrected using the Benjamini-Hochberg method [101], for decreasing the probability of making type I errors (false positives). From the average whole-brain model, comparisons between VGPs, NVGPs and CVGPs were performed using the Cohen’s D, for reporting the results in term of effect size. We considered this approach because *p*-values can be artificially inflated performing additional model realizations [102]. Effect sizes were defined as small for 0.2 < |D| < 0.5, moderate for 0.5 < |D| < 0.8, large for 0.8 < |D| < 1.2, and huge for |D| > 1.2.

### 4.13 Code and data availability

The scripts utilized for all simulations can be accessed via the following GitHub repository: https://github.com/carlosmig/StarCraft-2-Modeling.git. We used the Brain Connectivity Toolbox for Python (https://github.com/fiuneuro/brainconn) [47] for graph analysis. We obtained the Neurosynth association maps with the Python NiMARE Toolbox (https://nimare.readthedocs.io/en/latest/index.html) [50]. For generating the brain plots, we used the BrainNet Viewer Toolbox [103]. The structural connectivity data is obtainable upon request from the corresponding authors; however, a formal data sharing agreement must first be requested and signed. The empirical fMRI data was obtained from the Human Connectome Project Q3 release [39].

## Funding

CC-O is an Atlantic Fellow of the Global Brain Health Institute (GBHI) at Trinity College Dublin and is also supported by a postdoctoral grant from BrainLat. AI is partially supported by grants from ANID/FONDECYT Regular (1210195 and 1210176 and 1220995); ANID/FONDAP/15150012; ANID/PIA/ANILLOS ACT210096; FONDEF ID20I10152; ANID/FONDAP 15150012; and the MULTI-PARTNER CONSORTIUM TO EXPAND DEMENTIA RESEARCH IN LATIN AMERICA [ReDLat, supported by Fogarty International Center (FIC) and National Institutes of Health, National Institutes of Aging (R01 AG057234, R01 AG075775, R01 AG21051, CARDS-NIH), Alzheimer’s Association (SG-20-725707), Rainwater Charitable foundation – Tau Consortium, the Bluefield Project to Cure Frontotemporal Dementia, and Global Brain Health Institute)]. PO is partially supported by ANID/Fondecyt 1211750 and AC3E/Basal FB0008. This research was supported by the National Science Centre (Poland) Grant: 2013/11/N/HS6/01335, in the years 2013–2017 (to NK-G). The contents of this publication are solely the responsibility of the author and do not represent the official views of these institutions.

## Acknowledgments

The authors are grateful to Blizzard Entertainment Inc. for their help, and to StarCraft II players and controls who kindly participated in the experiment.

## Competing interests

None to declare.

## Author contributions

Conceptualization: CC-O, VM, PO, NK-G, AI.

Methodology: CC-O, VM, SO, JR, FL, JC, ET, PO, NK-G, AI.

Formal analysis: CC-O, VM, SO, JR.

Writing – original draft: CC-O, PO, NK-G, AI.

Writing – review & editing: CC-O, VM, SO, JR, FL, JC, ET, AB, PO, NK-G, AI.

Visualization: CC-O, VM, SO, JR.

Validation: CC-O, PO, NK-G, AI.

Data curation: CC-O, NK-G.

Supervision: ET, PO, NK-G, AI.

Project administration: AB, NK-G, PO, AI.

Resources: AB, PO, NK-G, AI.

## Notes

### Competing Interest Statement

The authors have declared no competing interest.

### Summary of Updates

New Results (Figure 8)

